# Unilateral Resection of Both Cortical Visual Pathways Alters Action but not Perception

**DOI:** 10.1101/2021.05.06.442988

**Authors:** Zoha Ahmad, Marlene Behrmann, Christina Patterson, Erez Freud

## Abstract

The human cortical visual system consists of two major pathways, a ventral pathway that subserves perception and a dorsal pathway that subserves visuomotor control. These pathways follow dissociable developmental trajectories, and, accordingly, might be differentially susceptible to neurodevelopmental disorders or injuries. Previous studies have found that children with cortical resections of the ventral visual pathway retain largely normal visuoperceptual abilities. Whether visually guided actions, supported by computations carried out by the dorsal pathway, follow a similar pattern remains unknown. To address this question, we examined visuoperceptual and visuomotor behaviors in a pediatric patient, TC, who underwent a cortical resection that included portions of the left ventral and dorsal pathways. We collected data when TC used her right and left hands to perceptually estimate the width blocks that varied in width and length, and, separately, to grasp the same blocks. TC’s perceptual estimation performance was comparable to that of controls, independent of the hand used. In contrast, relative to controls, she showed reduced visuomotor sensitivity to object shape and this was more evident when she grasped the objects with her contralesional right hand. These results provide evidence for a striking difference in the reorganization profiles of the two visual pathways. This difference supports the notion that the two pathways exhibit differential susceptibility to neurodevelopmental disorders.

## Introduction

The cortical visual system is comprised of two anatomically and functionally distinct pathways. The ventral pathway projects from the occipital lobe to the temporal lobe and supports vision-for-perception, while the dorsal pathway travels from the occipital lobe to the posterior parietal cortex and processes information that is utilized to support vision-for-action (Mishkin & Ungerleider, 1982; Goodale & Milner, 1992; for a revised view see Freud, Plaut & Behrmann, 2016; Freud, Behrmann & Snow, 2020).

The two pathways also differ in their maturation trajectories. Development of the dorsal pathway precedes that of the ventral pathway (Mundinano et al., 2015; Smith et al., 2017; Wattam-Bell et al., 2010; but see Ciesielski et al., 2019 for a different view). This earlier emergence relies, at least in part, on a transient pathway between the pulvinar and area MT (Bridge et al., 2016), which is necessary for the development of dorsal pathway structures and their associated behaviors (Kwan et al., 2021; Mundinano et al., 2018). Despite the earlier emergence, however, the dorsal pathway has a prolonged maturational trajectory, and functions associated with this pathway mature with or even after functions associated with the ventral pathway (Kiorpes et al., 2012). The combination of the early sensitivity and the prolonged developmental trajectory might give rise to the well-documented susceptibility of the dorsal pathway to neurodevelopmental disorders (Atkinson et al., 1997; Atkinson & Braddick, 2005; Atkinson, 2017).

In the current study, we sought to contrast directly the emergence of perceptual and visuomotor behaviors in a pediatric patient with a unique cortical resection that affected both pathways. Surgical resection of cortex can be effective in controlling seizures for individuals with pharmaco-resistant epilepsy. Temporal lobe surgical resection (one of the more common types of surgery given the site of the seizure focus) results in seizure remission for up to 80% of all patients, and long-term complete seizure freedom for up to roughly 41% of patients (Taylor et al., 2018). Depending on the nature of the underlying dysfunction, the surgery might entail a limited localized resection, a lobectomy or even functional disconnection or complete removal of a cerebral hemisphere (Lew, 2014). Pediatric patients with cortical resections provide a unique opportunity to advance the current understanding of the developmental trajectories of different cognitive functions as well as the nature and extent of cortical (re)organization and plasticity. Particularly, investigating the developmental patterns of behaviours mediated by specific cortical areas following the resection can elucidate the maturational chronology of the visual system. Thus, it is predicted that differential rates of maturation of visual cortical areas could elicit different trajectories for recovery of functions (Liu & Behrmann, 2017).

Previous research with post-surgery cortical resection patients has already reported on at least partial restoration of cognitive abilities such as intelligence (Skirrow et al., 2011; Vargha-Khadem et al., 1994), memory (Skirrow et al., 2015; Stretton et al., 2014), language (Ivanova et al., 2017; Nahum & Liegeois, 2020), and motor function (Gaberova et al., 2019; Jonas et al., 2004; McGovern et al., 2019). Recent studies that focus on perceptual behaviors have found that pediatric patients with resections that compromised a large portion of the ventral visual pathway typically demonstrate mostly normal visuoperceptual abilities. In particular, post-surgery visuoperceptual performance was found to be normal across a series of mid-level (for example, Glass patterns) and high-level (for example, face recognition) visual tasks (Liu et al., 2019). The normality in behavioral pattern was accompanied by normal topography, magnitude, and representational structure of category-selective organization in the non-lesioned hemisphere, as demonstrated using functional MRI. This conclusion was corroborated by a longitudinal study of a single child whose resection, at age 6 years and 9 months, resulted in the removal of the right occipital and posterior temporal lobes. Despite a persistent left homonymous hemianopia, the patient exhibited preserved intermediate- and high-level visual abilities suggesting a normal developmental trajectory following the resection (Liu et al., 2018).

Importantly, the preserved visuoperceptual behaviors described by these recent studies only examined computations carried out by the ventral visual pathway. However, the question remains whether visuomotor behaviors, mediated by the dorsal visual pathway (Goodale & Milner, 1992), display a similar pattern of resilience and follow a normal developmental trajectory post-resection. An alternative account is that the prolong developmental trajectory of the dorsal pathway (i.e., begins before and matures after the ventral pathway) may increase the susceptibility of visuomotor behaviors to early life injuries, including cortical resections.

To adjudicate between these alternatives, in the current study, we characterized the behavior of patient TC who had undergone a unilateral cortical resection that included portions of both the left ventral and dorsal pathways. We investigated her visuoperceptual and visuomotor competence using Efron blocks (Efron, 1969; Freud et al., 2016; Goodale et al., 1991). We expected that, consistent with the her intact performance reported before (Liu et al., 2019), TC would display normal perceptual abilities, presumably mediated by her intact right ventral pathway. In contrast, given the susceptibility of the dorsal pathway to early-life injuries, we predicted that TC’s visuomotor behaviors would be hindered, particularly when action engages the contralesional (right) hand.

## Methods

### Participants

TC, a right-handed 16-year-old female, and a control group of 14 neurotypical participants (10 female, average age 18.4 ± 1.6 years, all right-handed) were recruited for this study. TC suffered from epilepsy with an onset of seizures at the age of 7. Diagnosed with perinatal stroke with medically intractable focal epilepsy and multifocal encephalomalacia consistent with remote ischemic injury, she underwent surgery at the age of 13 years (Liu et al., 2019). Her surgery resulted in a left posterior parietal and occipitotemporal lobectomy (Figure 1). We delineated the extent of the resected region using a T1 MRI scan (resolution - 1mm^3^, Liu et al., 2019) obtained after the surgery and a detailed anatomical atlas (Mai, Majtanik & Paxinos, 2016). Close scrutiny of the anatomical scans revealed that most occipital structures were removed in the course of the surgery, including the posterior calcarine sulcus and Occipital gyri. Resected regions also include regions of the inferior temporal lobe (ventral pathway) such as the Fusiform and Lingual gyri. Additionally, the left Superior Temporal Sulcus (STS) is atrophied compared with the homologue right hemisphere sulcus. For the dorsal pathway, the resection includes posterior temporal cortex (adjacent to the proximate location of area MT) and posterior parietal cortex (i.e., Angular gyrus, posterior IPS). The more anterior portions of the intraparietal sulcus, known to be involved in visuomotor computations (Culham et al., 2003; Freud et al., 2018), are preserved.

**Figure 1:**
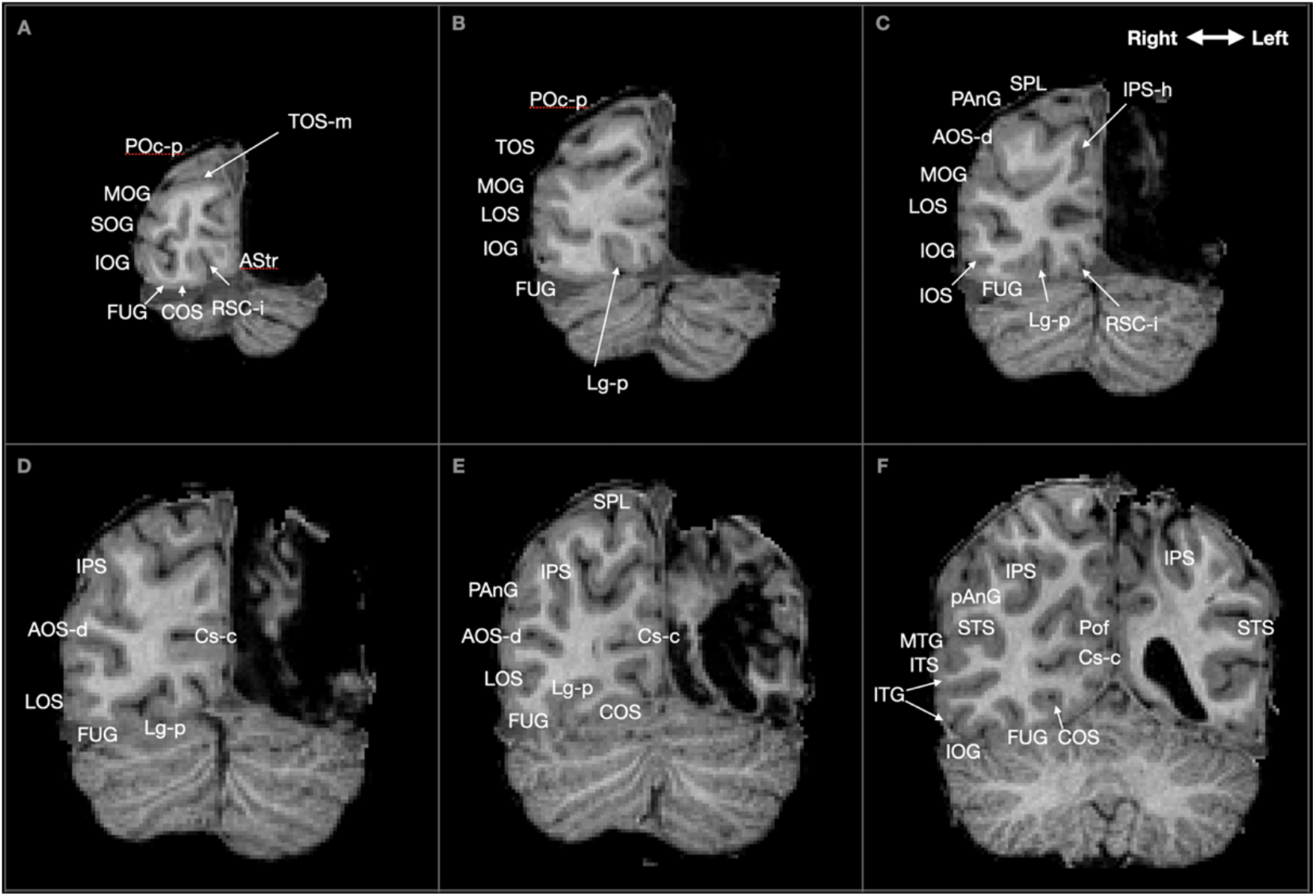
TC Postsurgical MRI (age 13 years). Representative coronal slices (posterior-A to anterior-F) from the MRI scan of TC. The resection included the posterior parts of the ventral and dorsal pathways of the left hemisphere. The regions homologous to the resected hemisphere were delineated using a detailed anatomical atlas (Mai, Majtanik & Paxinos, 2016). The resection extends to inferior and posterior parts of the temporal cortex. The resection includes all occipital structures, posterior temporal cortex (adjacent to the proximate location of area MT), and posterior parietal cortex (i.e., angular gyrus, posterior IPS). Identified areas include the anterior occipital sulcus, dorsal segment (**AOS-d**); striate area (**AStr**); collateral sulcus (**COS**); calcarine sulcus (**Cs-c**); fusiform gyrus (**FUG**); inferior occipital gyrus (**IOG**); inferior occipital sulcus (**IOS**); intraparietal sulcus (**IPS**); intraparietal sulcus, horizontal segment (**IPS-h**); inferior temporal gyrus (**ITG**); inferior temporal sulcus (**ITS**); lingual sulcus, posterior ramus (**Lg-p**); lateral occipital sulcus (**LOS**); middle occipital gyrus (**MOG**); middle temporal gyrus (**MTG**); posterior angular gyrus (**PAnG**); posterior-occipital arc, posterior part (**POc-p**); parietooccipital fissure (**Pof**); retrocalcarine sulcus, inferior branch (**RSC-i**); superior occipital gyrus (**SOG**); superior parietal lobule (**SPL**); superior temporal sulcus (**STS**); transverse occipital sulcus (**TOS**); transverse occipital sulcus, medial ramus (**TOS-m**).

For the current study, TC was tested at her home and provided assent, and her parents provided informed consent for her participation. Control participants were tested using the same experimental setup (see Apparatus and Stimuli for details) at York University, Toronto. Participants older than 18 years of age provided informed consent. Minor participants provided assent and their parents provided informed consent. Participants received course credit or $15 as compensation for their participation. The experimental protocol was approved by the Institutional Review Board of Carnegie Mellon University and by York University Human Participants Review Committee.

### Data availability

Raw data as well as the analysis code are distributed under the terms of the Creative Commons Attribution License, which permits unrestricted use and redistribution provided that the original author and source are credited. https://osf.io/c4qky/?view_only=91dcd53067284a298ee7b9a056532f06

### Apparatus and Stimuli

Participants sat in front of a table on which the target objects were presented. The target objects were a set of four Efron blocks (1969) that all had the same surface area, texture, and color, but varied in width and length. The width of the blocks ranged from 20 - 35 mm in gaps of 5mm and lengths adjusted accordingly (see Figure 2A). Grasping movements and manual estimations were recorded using an Optitrack system (Natural Point DBA OptiTrack, USA). The system included four Prime 13W cameras and three active infra-red-light emitting diodes attached to the participant’s hand in such a way that permitted complete freedom of movement of the hand and fingers (Figure 2B). The system tracked the 3D trajectory of the participants’ index, thumb and wrist movement using a 100 Hz sampling rate and allows to calculate the aperture between the fingers at any given time point.

**Figure 2:**
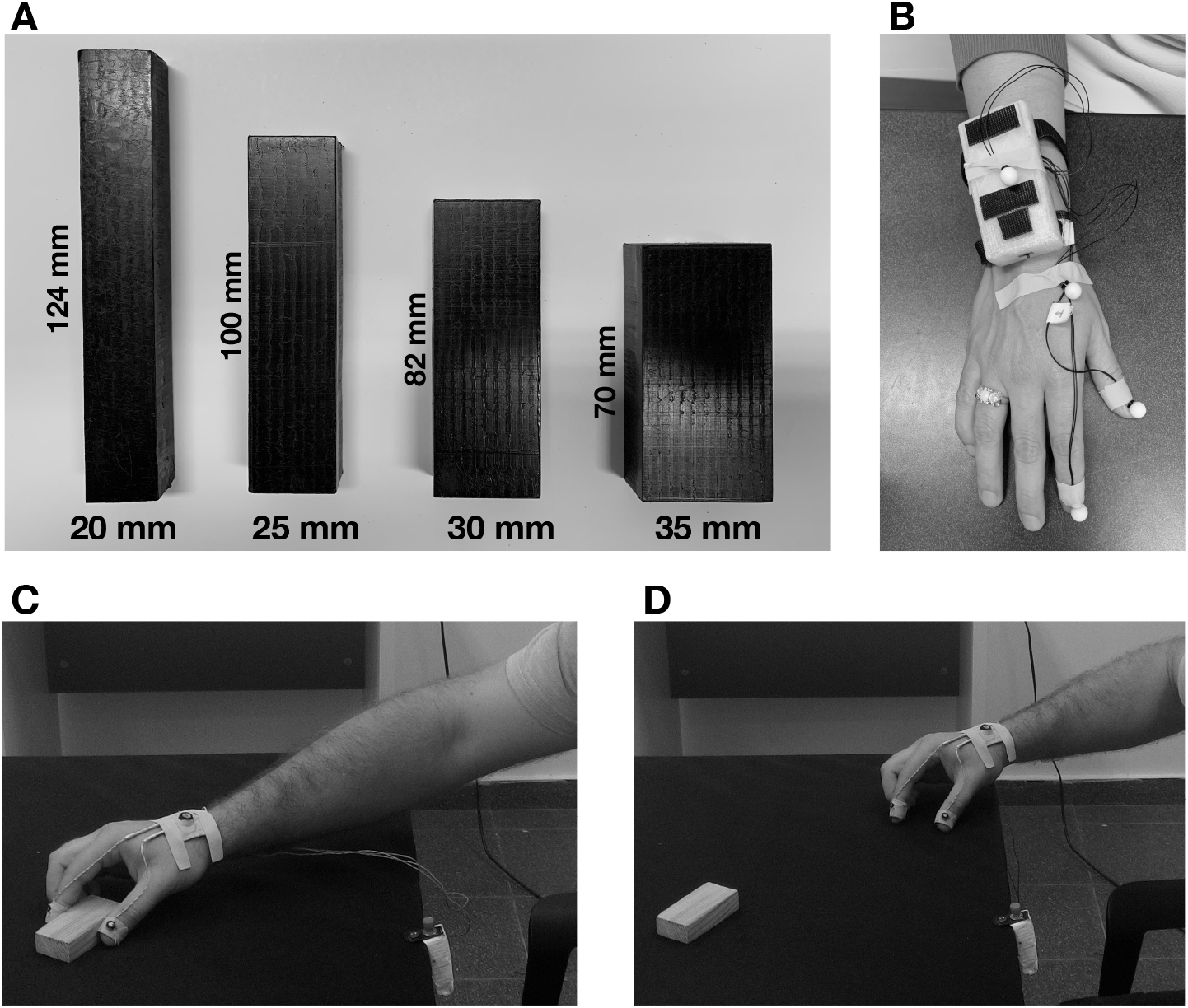
**(A)** Experimental stimuli - target objects used in the experimental set-up were a set of four Efron blocks that all had the same surface area, texture, mass, and color, but varied in width and length, with their width indicated below the block. Using their index finger and thumb, in separate blocks of trials, participants were asked either to grasp the blocks or manually estimate the width of the blocks. **(B)** Location of diodes **-** three active infra-red-light emitting diodes were attached to the participant’s hand during the experiment for tracking the grasping and estimation trajectories. **(C)** Grasping trial – example of grasping trial, in which the participant reached and grasped one of the target objects **(D)** Manual estimation trial - example of manual estimation trial, in which the participant indicated the width of the target objects with their thumb and index finger.

### Procedure

Participants completed two tasks, a grasping (Figure 2C) and manual estimation task (Figure 2D). In each task, participants began with their thumb and index finger grasping a permanently stationary block immediately in front of them. This was referred to as the “home” position. On each trial, one of the four target objects was placed in front of the participant with the width parallel to the left-right orientation of participant, within arm’s reach (approximately 40 cm). In the grasping task, the participants were required to reach for the target object with the thumb and index finger across its width (thumb more proximal to viewer and index finger more distal) and to lift it off the table (approximately 15 cm) before setting it down and then returning to the “home” position. In the manual estimation task, participants were required to indicate the perceived width of object by extending their thumb and index finger at a height of approximately 15 cm from the table surface to estimate the corresponding width. They were instructed to hold the finger posture for two seconds before returning to the “home” position. Each task was completed separately with each hand, resulting in four experimental blocks. In each block, each of the four target objects was presented 15 times in a randomized order resulting in a total of 60 trials per block. All participants completed the experiment in the following order: grasping using the right hand, manual estimation using the right hand, grasping using the left hand, manual estimation using the left hand to mirror the same order as used for TC.

### Data analysis

For each trial, the 3D trajectory of the index finger and thumb was analyzed using in-house code written in Python. The starting point of the grasping movement was defined as the frame following five consecutive frames that had a velocity greater than 10mm/sec. The endpoint of the grasping movement was defined as the point during three consecutive frames in which the change in grasping aperture (i.e., the distance between the thumb and the index finger) relative to the previous frame was smaller than 0.2mm. An additional condition was that the Z (superior-inferior) location of the fingers was smaller than 80mm, which indicated that the fingers were positioned along the same plane as the target object. The Maximum Grip Aperture (MGA) was calculated for each trial as the frame that reached the maximum distance between the index finger and the thumb between the time after the movement onset and the end of the movement. For the estimation task, the aperture between the thumb and index finger that was held constant over 10 consecutive frames was determined to reflect the perceived width of the object (Freud et al., 2016). All trials were visually inspected, and the analysis was manually refined for a small number of trials in which the algorithm did not accurately detect the end point of the movement.

The Just Noticeable Difference score (JND) and correlation between the MGA and object width (Fisher transformed) were calculated for each participant separately for each task. The JNDs were measured by analyzing the standard deviation in the MGA for each object in each task (Freud et al., 2016; Ganel et al., 2008). The JND measures the minimum detectable increment in stimulus magnitude and therefore reflects the sensitivity, which is the size resolution in this case, of the task of interest (Marks & Algom, 1998). Pearson’s correlation (which was subsequently transformed to a Fisher’s z-score to allow statistical analysis) was determined by correlating the average MGA for each target object with the real size of the object. Previous research has indicated that a high correlation between the MGA and the real object size is a strong reflection of the integrated relationship between the physical demands of a grasping task and the resulting fine motor control in response to those demands ( Goodale et al., 1991).

### Grasping Similarity Analysis

In addition to the analyses described above, we employed a grasping similarity analysis (GSA) in order to characterize visuomotor sensitivity along the entire motion trajectory and to permit a more fine-grained multi-dimensional description of shape sensitivity. This approach is based on representational similarity analysis (Kriegeskorte et al., 2008), often used in computational neuroscience. In this method, representational dissimilarity matrices (RDMs) are computed to characterize the information carried by a given representation in a model. Here, we utilized this approach to compare the similarity of the grasping trajectories directed to the different objects. Importantly, in contrast to the previous analyses mentioned above, this approach does not depend on the MGA, but rather takes into account the entirety of the motion trajectory.

The analysis was done separately for each participant on the grasping data. For each trial, we extracted the grip aperture along the movement trajectory and correlated it with the grip aperture of all other trials of all sizes (Figure 3A). Notably, in order to estimate the similarity between trials with different lengths (i.e., movement times), we applied Dynamic Time Warping (DTW) which calculates the distance between two time series of different lengths (Giorgino, 2009) (Figure 3B). For the estimation task, similar to previous studies (e.g., Freud et al., 2016; Ganel et al., 2008; Goodale et al., 1991), only the final aperture was used to analyze sensitivity to object size and therefore GSA analysis was not employed.

**Figure 3:**
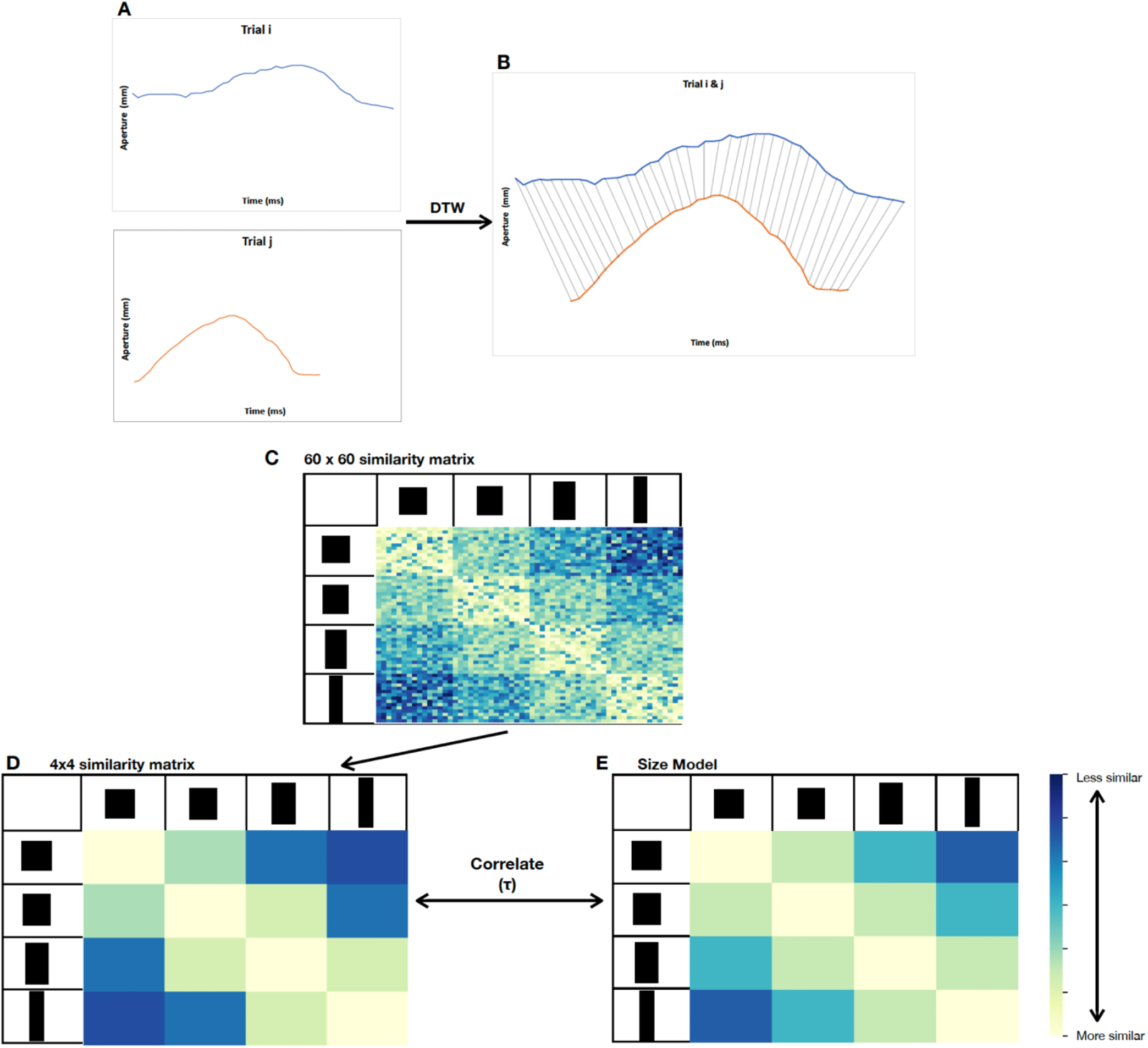
GSA methodology. **(A)** The grip aperture along the movement trajectory was extracted for each participant and each trial. **(B)** Dynamic Time Warping was applied in order to estimate the similarity between each pair of trials. **(C)** This process yielded a 60 × 60 similarity matrix (per subject, per run; 15 trials per stimulus x 4 size). **(D)** The 60 × 60 similarity matrix was used to create a 4×4 similarity matrix by averaging the distances per size. **(E)** The 4×4 similarity matrix was correlated with a hypothetical size model which reflected perfect sensitivity to size information. The Kendall rank correlation coefficient between the model and the observed similarity matrices was computed separately for each hand of each participant.

This analysis yielded a 60 by 60 dissimilarity matrix per run (15 trials per stimulus X 4 size) (Figure 3C), where lower values reflect greater similarity in movement trajectories of any given pair of trials. Next, we averaged the distances per size resulting in a 4X4 asymmetrical matrix (Figure 3D). Finally, we defined a model that reflects perfect sensitivity to size information such that objects that are more similar in size (e.g., 20mm vs 25mm) are expected to elicit more similar grasping trajectories compared with objects that are dissimilar in their graspable size (e.g., 20mm vs 35mm) (Figure 3E). We then computed the Kendall rank correlation coefficient between the model and the observed similarity matrices. It was anticipated that TC would have grasping RDMs that deviate from the size model more so than those of the controls due to deficits in visuomotor sensitivity following her resection.

Previous research has indicated that the movement trajectory analysis might produce false sensitivity to object size, particularly for the closing (i.e., post-MGA) portion of the grasping movement due to time-normalized data (Whitwell & Goodale, 2013). Hence, we repeated the RSA procedure described above, but defined the MGA of each trial as the endpoint of this trial and the results reported below (see results section) were fully replicated for this revised analysis. Thus, sensitivity to object size observed using the GSA analysis could not be attributed to problem in the normalization of the grasping trajectory.

### Statistical analysis

We applied a modified single-subject t-test to examine whether TC’s scores across the different variables deviated from the performance of the control groups across the different tasks and dependent measures (see above) (Crawford & Garthwaite, 2005). Finally, we also used the Revised Standardized Difference Test (RSDT) (Crawford & Garthwaite, 2005) to measure whether the difference between TC’s standardized score on two conditions (e.g., grasping with the right hand and grasping with the left hand) was significantly different from the difference measured in the control sample.

## Results

To examine whether TC was impaired in the perceptual and/or grasping task, we examined her performance in the two tasks, each completed with her ipsilesional (left) and contralesional (right) hands.

### Average aperture

First, we analyzed the average aperture across the different object sizes for each task. Interestingly, TC exhibited final apertures that fell within the normal range for the manual estimation task (Figure 4A, left), as verified by single-case statistical comparisons [right hand: t_(13)_ = 1.63, p>.1, 1.68 (0.84 to 2.05); left hand: t_(13)_ < 1, 0.41 (−0.14 to 0.95)]. However, for the grasping task, her maximum grip apertures for both her contralesional and ipsilesional hands that fell outside of the normal range (Figure 4A, right, grasping left: 92.11 mm, grasping right: 96.16 mm). In fact, her MGA was, on average, ~20mm larger compared with that of control participants, and, single-case statistical comparisons confirmed this exaggeration for both the right [t_(13)_ = 3.24, p < 0.05, Z-_CC_ = −2.099 (1.97 to 4.73)] and left hand [t_(13)_ = 2.816, p < 0.05, Z-_CC_ = 2.91 (1.68 to 4.12)]. This finding is consistent with previous reports that demonstrated that disproportionately large aperture is indicative of a visuomotor deficit (Jakobson et al., 1991)

**Figure 4:**
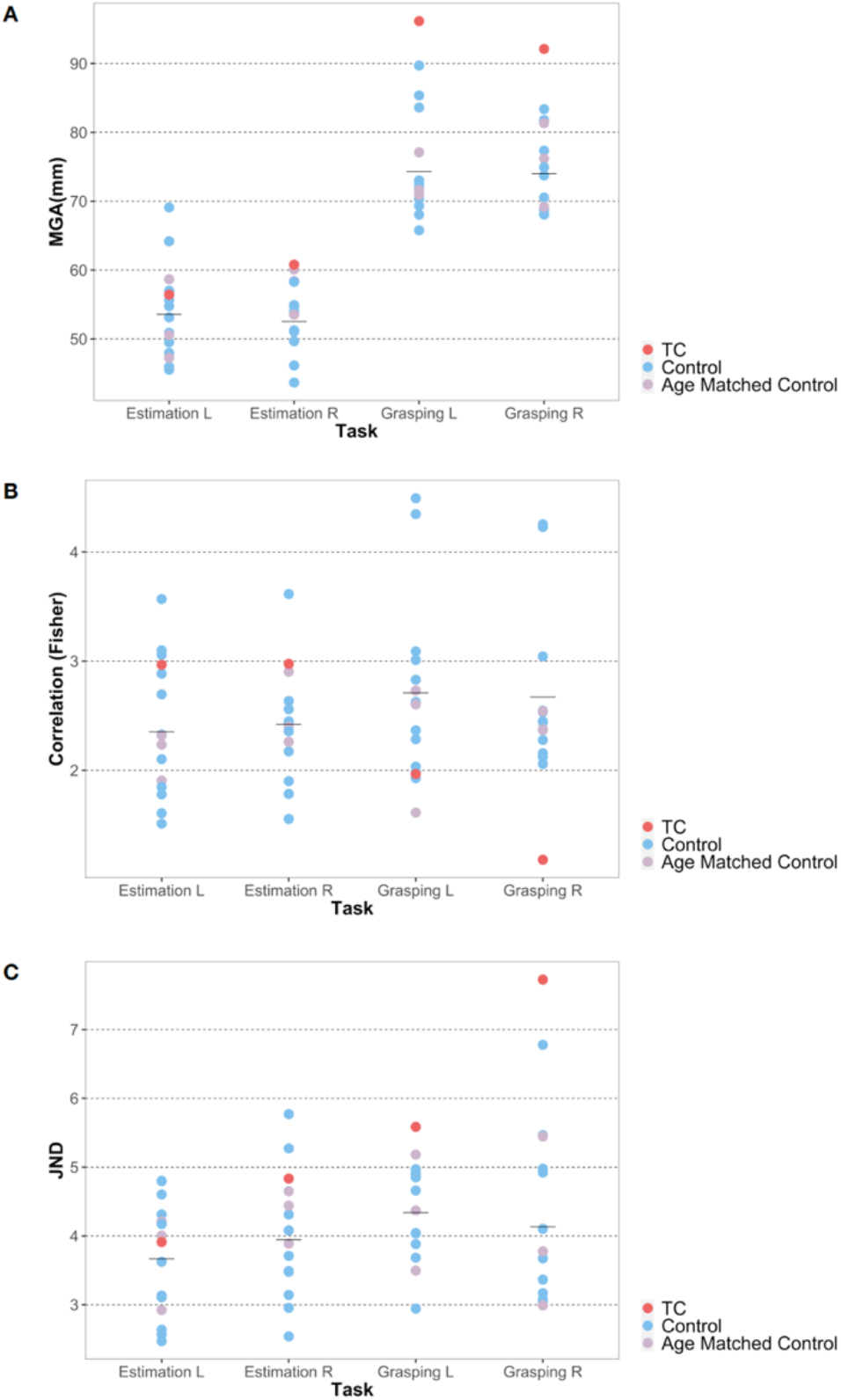
Results from grasping and manual estimation tasks. Across all figures, TC’s value is represented as the dark red dot. Each blue (control) or purple (age-matched control) dot represents the value of a single participant’s data. The mean value of the control group is indicated by the horizontal black line. R stands for right hand and L stands for left hand. **(A)** The average results of the MGA (grasping) or final aperture (estimation) in mm for each task. TC exhibited normal final apertures for the manual estimation task but showed exaggerated maximum grip apertures for the grasping task. **(B)** The correlation results between the true object size and hand aperture for all tasks. Higher values indicate greater sensitivity. TC showed lower correlation than in the controls just for the grasping task and only in her right hand. **(C)** The JND values representing the average within-subject variability to each Efron block. Higher values represent reduced sensitivity to object size. TC was found to have impaired resolution in the grasping task, but not in the manual estimation task.

Next, using the RSDT (Crawford & Garthwaite, 2005) to determine whether the difference between TC’s grasping performance was different between her two hands, we observed no difference between her left and right hand [t_(13)_< 1, Z-DCC = −0.238 (95% CI = −1.780 to 1.266)].

### Size – hand aperture correlation

Next, to characterize sensitivity to object shape, we computed the correlation between grasping and estimation aperture to object size where a correlation value of 1 indicates an ideal fit between object width and hand aperture. For control participants, we found a high correlation between object size and hand aperture across hands and tasks (Figure 4B). In accordance with previous research (Liu et al., 2019), for the perceptual task, we found a comparable sensitivity to that of controls regardless of hand used by TC [right hand t_(13)_ = 1.04, p > 0.2, one-tailed, Z-_CC_ = 1.077 (95% CI = 0.399 to 1.729); left hand [t_(13)_ = 0.093, one-tailed, Z-_CC_ = 0.097 (95% CI = −0.430 to 0.620)]. In contrast, TC’s sensitivity to object size was significantly impaired for the grasping task but only with her right (contralesional) hand [t_(13)_ =−2.027, p < 0.05 one-tailed, Z-_CC_ = −2.099 (95% CI = −3.041 to −1.133)] and not with the left (ipsilesional) hand, with TC’s correlation values falling within the normal range [t_(13)_ = −0.851, one-tailed, Z-_CC_ = −0.881 (95% CI = −1.491 to −0.247)].

Next, we employed the RSDT (Crawford & Garthwaite, 2005) to statically validate the dissociation between grasping and estimation performance for her right hand [t_(13)_ = 1.849, p < 0.05, one-tailed, Z-_DCC_ = 1.979 (95% CI = 1.081 to 2.957)]. In contrast, for the left hand, there was no difference between the two tasks [t_(13)_ < 1, Z-_DCC_ = 0.768 (95% CI = 0.162 to 1.408)]. Finally, we did not find a significant difference between TC’s performance for grasping with her right hand compared to her left hand [t_(13)_ = 1.039, p > 0.2, Z-DCC = −1.140 (95% CI = −2.151 to −0.236)].

### JNDs

The average within-subject variability of responses to each Efron block was used as an additional indicator of sensitivity to the objects’ width (Freud et al., 2016; Ganel et al., 2008). Here, smaller values reflect finer resolution for object size.

Consistent with the correlation results (see above), analysis of the JND values indicated that TC’s performance for the manual estimation tasks was comparable to the control group mean, confirming that she exhibited normal sensitivity to object shape in the manual estimation task for both her right hand [t_(13)_ < 1, Z-_CC_ = −0.247 (−0.775 to 0.290)] and left hand [t_(13)_ < 1, Z-_CC_ = 0.273 (−0.266 to 0.802)] (Figure 4C, left). TC’s variability was greater than that of controls for the grasping task (Figure 4C, right) for both her right (contralesional) [t_(13)_ = 2.923, p < 0.01 one-tailed, Z-_CC_ = −2.099 (−3.041 to −1.133)] and her left (ipsilesional) hand [t_(13)_ = 1.7891, p < 0.048 one-tailed, Z-_CC_ = 1.851 (0.961 to 2.715)].

The RSDT test confirmed the existence of a dissociation between grasping and estimation performance with the right hand [t_(13)_ = 2.272, p < 0.03, one-tailed, Z-_DCC_ = 2.472 (95% CI = 1.346 to 3.785)]. In contrast, no significant difference between the perceptual and action tasks was observed for the left hand [t_(13)_ = 1.40, p > 0.05, one-tailed, Z-_DCC_ = −1.121 (95% CI = −1.946 to 0.376)]. Finally, the RSDT (Crawford & Garthwaite, 2005) did not provide evidence for a dissociation between TC’s performance for grasping with her right compared to her left hand [t_(13)_ = < 1, Z-DCC = 0.861 (95% CI = −0.225 to 2.061)].

### Grasping Similarity Analysis of grasping kinematics

Next, we utilized a novel GSA approach to uncover the similarity across grasping trajectories directed to different objects. It is expected that, for the control participants, objects with widths similar to one another will have a more similar movement trajectory (e.g., the 20mm and 25 mm Efron blocks) while objects that differ the most from each other (e.g., the 20mm and 35 mm Efron blocks) will have the most dissimilar trajectory.

Typical observers’ grasping trajectories showed high correspondence with the size model for both the right and left hand (Figure 5A, 5B), with the average Tau scores well above zero (right hand [t_(13)_ = 41.58, p < 0.0001, Cohen’s *d* = 11.11], left hand [t_(13)_ = 41.79, p < 0.0001, Cohen’s *d* = 11.17]). This high correspondence indicates that the shape model explains the movement trajectory data well and validates this analytical approach.

**Figure 5:**
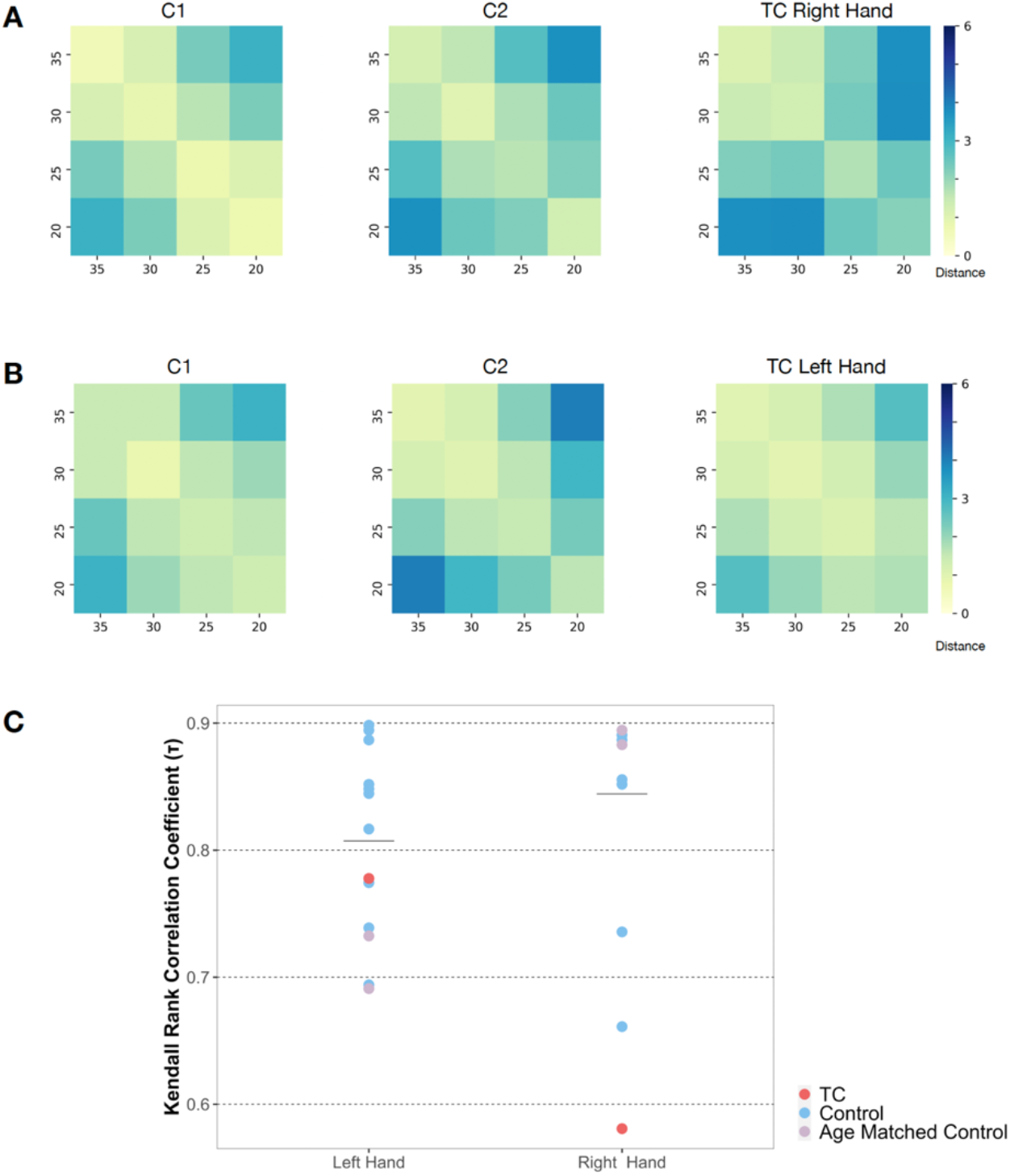
GSA results. **(A)** GSA matrices for TC’s right hand and two representative control participants. Lower distance (bright colors) values reflect greater similarities between trials. Each Efron block width is indicated by the ticks on the axes of the matrices (in mm). **(B)** GSA matrices for TC’s right hand and two representative control participants. (**C)** Each participant’s Kendall rank correlation coefficient is reported for both hands. TC is significantly different than the controls for the grasping task in her right (contralesional) hand. TC is comparable to the controls for the grasping task in her left (ipsilesional) hand.

Next, we compared the Tau value observed for TC with that observed for the controls. For grasping with her left hand, TC exhibited similar correspondence with the size model compare to that of the controls [t_(13)_ < 1, Z-_CC_ = −0.429 (−0.970 to 0.127)] (Figure 5C, left). In contrast, when grasping with her right hand, TC’s grasping trajectories were significantly less correlated with the size model compared with controls [t_(13)_ = −3.140, p < 0.008, Z-_CC_ = −3.250 (−4.581 to −1.900)] (Figure 5C, right). These results suggest that TC’s visuomotor impairment is not limited to her MGA results but is also evident along the movement trajectory as a whole (as she is not scaling her hand appropriately for the objects).

## Discussion

The current study was designed to elucidate possible dissociable effects of a unilateral cortical lesion of the dorsal and ventral pathways on visuomotor and perceptual behaviors. We examined shape sensitivity in TC, an adolescent who has a left lateralized cortical resection that affects both visual pathways. Notwithstanding the ventral resection, TC displayed preserved perceptual sensitivity to object shape. In contrast, her visuomotor sensitivity was profoundly impaired when she used her right, contralesional hand and, to a lesser extent, when she used her left, ipsilesional, hand. This deficit was observed across different dependent measures including aperture size, variability of the grasping aperture and sensitivity to object size (see Table 1 for a summary of TC’s performance).

**Table 1:** Summary of TC’s performance on behavioural tasks compared to controls.

| Measure | Task (Hand) |  |  |  |
| --- | --- | --- | --- | --- |
|  | Grasp |  | Estimation |  |
|  | LH | RH | LH | RH |
| Average aperture | × | × | ✓ | ✓ |
| Size – hand aperture correlation | ✓ | × | ✓ | ✓ |
| Just Noticeable Difference | × | × | ✓ | ✓ |
| Grasping Similarity Analysis | ✓ | × | N/A | N/A |

Note: Checks (✓) indicate TC's performance was comparable to that of the controls for that measure. Impairment of each hand (left/right) for each task (grasp/estimate) is indicated by crosses (×).

The results from the estimation tasks are consistent with previous investigations that documented retained perceptual functions in pediatric patients with cortical resections even when the resection compromised a large portion of the ventral visual pathway (Granovetter et al., 2020; Liu et al., 2019). Our results revealed a consistently different pattern of behavior for the visuomotor task, presumably mediated by computations carried out by the dorsal pathway. Hence, the current study provides evidence for dissociable post-surgical profiles of behaviors associated with the two visual pathways.

The reasons for the dissociable effects of the cortical resection on visuomotor and perceptual behaviors are not entirely clear. One explanation might be that TC’s dorsal lesion was more extensive and included more critical regions for visuomotor control. Note, however, that careful delineation of the lesion does not support this explanation (Figure 1). In particular, multiple structures along both pathways were affected. Notably, for the ventral pathway, the lateral occipital cortex, which is known to be critical for shape perception (Grill-Spector et al., 2001), was resected, while the anterior portion of the left IPS, which plays a critical role in visuomotor control (e.g., Culham et al., 2003; Freud et al., 2018) was not resected. Based on those observations, we propose that the dissociation between perception and action is more likely to reflect the differential maturation rates of the two visual pathways.

Previous studies have shown that the dorsal pathway begins to develop before the ventral pathway (Mundinano et al., 2015; Wattam-Bell et al., 2010) but is then subject to a prolonged developmental trajectory (Braddick et al., 2003; Kiorpes et al., 2012). Moreover, the normal development of the dorsal pathway, and, accordingly, of visuomotor behaviors, depends, at least partially, on a transient pathway from the pulvinar to area MT that projects to the parietal cortex (Kwan et al., 2021; Mundinano et al., 2018). These unique properties of dorsal pathway development might give rise to well-documented sensitivity of this pathway to neurodevelopmental disorders (Atkinson, 2017, Atkinson & Braddick, 2005; Braddick et al., 2003), and can also account for the results reported here.

It is worth noting that the proximal location of area MT that is critical for the development of visuomotor control (Kwan et al., 2021; Mundinano et al., 2018) was partially resected in TC. Given that TC suffered from an early stroke, it is possible that this region was comprised early in life and the observed visuomotor deficits specifically reflect that lack of necessary input from this region to parietal structures. This question might be partially addressed in future studies with patient TC that will utilize neuroimaging tools to characterize the functional and connectivity properties of area MT in the two hemispheres.

### Bilateral deficit after a unilateral lesion

TC’s resection is confined to the left hemisphere. However, despite this clear laterality, her visuomotor deficit is also evident when she grasped with her left, ipsilesional hand, albeit to a lesser extent. There are two possible neural mechanisms, which are not mutually exclusive, that might account for the bilateral nature of TC’s visuomotor deficit, namely hemispheric specialization within the dorsal pathway or an inter-hemispheric inhibition process.

Hemispheric specialization refers to the dissociable contribution of each hemisphere to different functions. This specialization is not strictly dichotomous, but is reflected on a continuum of functions between the hemispheres (Bradshaw & Nettleton, 1981). For example, in most people, both right and left handed, language is more lateralized to the left hemisphere (Knecht et al., 2000; Ojemann, 1991), despite a contribution of the right hemisphere to different aspects of language (Ross & Mesulam, 1979; Vigneau et al., 2011).

The notion of hemispheric specialization has also been demonstrated for the dorsal visual pathway (i.e., parietal cortex) such that while the left hemisphere plays a major role in visuomotor computations even among left handed individuals (Gallivan & Culham, 2015; Gonzalez et al., 2006), the right hemisphere contributes to attentional mechanisms (Bowen et al., 1999; Ringman et al., 2004; Becker & Karnath, 2007) and spatial transformations (Gauthier et al., 2002.; Harris et al., 2000; Warrington & Taylor, 1973).

This specialization is also supported by neuropsychological investigations. For example, patients with right-hemisphere injuries displayed a preserved dissociation between action and perception, which is not evident for patients with left-hemisphere lesions (Radoeva et al., 2005). Additionally, greater severity of optic ataxia (Perenin & Vighetto, 1988) was observed in patients with left hemisphere injures. In particular, most patients with optic ataxia after a left hemisphere lesion have displayed a hand effect (errors when pointing with their contralesional hand) as well as a contralateral field effect (errors when pointing to stimuli in the contralesional visual hemifield) (Vindras et al., 2016), whereas patients with a right hemisphere lesion showed a milder version of optic ataxia with only a field effect (Perenin & Vighetto, 1988). TC’s deficit is consistent with the non-dichotomous specialization of the left hemisphere in visuomotor computations. In particular, despite the unilateral nature of her lesion, her grasping behaviors were altered when she used her right (contralesional) and, albeit to a lesser extent, when she used her left (ipsilesional) hand.

A second possible process that could have contributed to this bilateral decrement is inter-hemispheric inhibition of the non-lesioned right parietal cortex. Inter-hemispheric inhibition refers to the process by which one perturbed hemisphere of the brain inhibits the opposite hemisphere (van Meer et al., 2010). This phenomenon was described in a case of visual agnosia after a lesion sustained to the right ventral pathway. Despite the unilateral nature of the lesion, reduced visual responses, object-related and -selective responses were also observed in homologous locations in the structurally intact left hemisphere, pointing to a diaschisis of regions in the non-lesioned hemisphere (Konen et al., 2011; Freud & Behrmann, 2020). Importantly, inter-hemispheric inhibition was also described in the context of motor behaviors (Murase et al., 2004), and it was demonstrated that reducing this inhibition using TMS can contribute to motor training (Williams et al., 2010).

The bilateral nature of TC’s deficit is consistent with the inter-hemispheric inhibition account, such that the left lesion adversely affected activation in the non-lesioned, right parietal cortex. In contrast, it is not clear why inter-hemispheric inhibition would affect only one pathway and not the other. Thus, to test this hypothesis, future studies, with cortical resection patients, should utilize a neuroimaging approach to better describe visuomotor and perceptual representations across the two hemispheres and to evaluate the connectivity patterns.

### GSA as a tool for the investigation of size resolution in grasping tasks

Previous studies that have utilized the Efron task to investigate sensitivity to object shape, did so by calculating the correlation between the hand aperture and the object width (Goodale et al., 1991; Karnath et al., 2009). Some later studies also used JNDs based on the within-subject standard deviations of the hand aperture as an additional dependent measure to investigate the resolution of the scaling of hand aperture to object width (Freud al., 2016). Note that both of these measures are focused on a single point taken from the full movement trajectory (the MGA) and, therefore, do not capture the details of the movement trajectory as a whole.

In the present investigation, we further extended the analyses that measure the resolution of response in grasping tasks by adjusting the well-established representational similarity analysis (RSA) (Kriegeskorte et al., 2008) to measure similarity of grasping trajectories directed to different objects. Importantly, this approach takes into account the entire course of the motion trajectory. Moreover, this analysis allowed us to describe shape sensitivity in a multidimensional space (i.e., similarity of movements directed to different object size) and to correlate directly theoretical, ideal, models with the observed data.

As expected, the GSA revealed a strong correspondence between grasping similarity matrices and the predicted size model for control participants. Consistent with the results obtained for the MGA analysis, TC’s profile showed a poor correspondence between grasping similarity and the size model only for grasping movements completed with her right contralesional hand. These results indicate that TCs’ profound visuomotor impairment was not limited to her MGA results but was evident along the movement trajectory as a whole. Moreover, these results provide a proof-of-concept that the GSA approach has utility as an additional measure of sensitivity to objects’ dimensions in grasping tasks in typical and atypical populations.

### Limitations

The current study provides important insights into the effect of a unilateral cortical resection on visuomotor and visuoperceptual behaviours. However, several limitations should be noted and perhaps addressed in future experiments.

First, we note that TC suffered from three related neurological incidents. She was diagnosed with perinatal stroke and then suffered from epilepsy with an onset of seizures at the age of seven. Diagnosed with medically intractable focal epilepsy and multifocal encephalomalacia consistent with remote ischemic injury, she had a cortical resection at the age of 13 years. As such there is no concrete way of knowing to what degree each of these incidents resulted in her performance in the current study.

Second, the current study is based solely on TC’s behavioral performance. As such, it is impossible to conclude whether the retained perceptual behaviors rely on the intact right hemisphere, or alternatively on remaining tissue in the left hemisphere. Importantly, however previous investigation of TC’s neural profile confirmed that her affected left hemisphere ventral pathway was not sensitive to any of the tested visual categories (faces, objects, words and scenes), while normal sensitivity was observed along the right ventral pathway (Liu et al., 2019). Hence, it is reasonable to assume that the retained perceptual abilities observed in this patient were mediated by computations carried out by the intact right occipitotemporal cortex.

## Conclusion

The goal of the current study was to explore the effect of an early onset unilateral lesion affecting both visual pathways on perception and action. We found that perceptual behaviours presumably mediated by the ventral pathway were retained, while visuomotor behaviours presumably mediated by the dorsal pathway were selectively impaired. These results provide novel evidence for fundamental differences in the reorganization profiles of the two visual pathways, which might reflect the differential developmental trajectories of visuomotor and perceptual behaviors.

## Acknowledgements

This research was supported by the Natural Sciences and Engineering Research Council of Canada (NSERC) (EF), by the Vision Science to Applications (VISTA) program funded by the Canada First Research Excellence Fund (CFREF, 2016–2023) (EF) and by RO1 EY027018 from the National Institutes of Health to MB and CP. The authors would like to thank TC and family for their participation and interest.

